# Geographic drivers of diversification in loliginid squids with an emphasis on the western Atlantic species

**DOI:** 10.1101/2020.07.20.211896

**Authors:** Gabrielle Genty, Carlos J Pardo-De la Hoz, Paola Montoya, Elena A. Ritschard

**Affiliations:** Departamento de Ciencias Biológicas, Universidad de los Andes, Bogotá D.C, Colombia; Department of Biology, Duke University, Durham, North Carolina, 27708, United States of America; Instituto de Investigación de Recursos Biológicos Alexander von Humboldt, Bogotá, D.C., Colombia; Department of Neuroscience and Developmental Biology, University of Vienna, Austria

**Author notes:** These authors contributed equally to this work (Gabrielle Genty, Carlos J Pardo-De la Hoz, Elena A. Ritschard). Correspondence author: Gabrielle Genty.

**Keywords:** Loliginidae, Historical biogeography, ecological isolation, cryptic speciation

## Abstract

**Aim:** Identifying the mechanisms driving divergence in marine organisms is challenging as opportunities for allopatric isolation are less conspicuous than in terrestrial ecosystems. Here, we aim to estimate a dated phylogeny of the squid family Loliginidae, and perform ecological niche analyses to explore biogeographic and evolutionary patterns giving rise to extant lineages in this group, with particular focus on cryptic species with population structure along the western Atlantic coast.

**Location:** World-wide.

**Taxon:** Class Cephalopoda, Family Loliginidae

**Methods:** We used three loci to infer gene trees and perform species delimitation analysis to detect putative cryptic speciation events. We then estimated a dated species tree under the Bayesian multispecies coalescent and used it to reconstruct ancestral distributions based on the currently known ranges of the species. Also, we tested the hypothesis of niche divergence in three recently diverged species subpopulations of the northwestern and southwestern Atlantic Ocean by ecological niche modeling and niche overlap measurement from occurrence data.

**Results:** The phylogenetic analyses confirmed the monophyly for the current twenty-six species of the Loliginidae family. Our ancestral area reconstruction and divergence estimation revealed the origin and geographical dispersal of loliginid lineages. Additionally, the phylogenetic analysis and the species delimitation analysis supported geographic structure within *D. pleii, D. pealeii* and *L. brevis.* The ecological niche models revealed unsuitable habitat in the immediately adjacent area of the Amazonian Orinoco Plume, yet suitable habitat characteristics beyond this area.

**Main conclusions:** Our study allowed us to confirm the monophyly of all currently recognized species within the Loliginidae family and we corroborate the biogeographical origin being the Indo-Pacific region in the Cretaceous. We found a possible new cryptic lineage and show evidence of the Amazon-Orinoco Plume as an ecological barrier, which influenced the diversification of this particular group of marine organisms.

## INTRODUCTION

Geographic divergence can be seen as a continuum of processes causing isolation via dispersal barriers and ecological gradients varying across biogeographic regions (Cowman & Bellwood, 2013; Pyron & Burbrink, 2010). Hard physical barriers causing vicariance are often thought to be the major force driving divergence and the consequent speciation processes in terrestrial ecosystems (Coyne & Orr, 2004; Smith et al., 2014; Wiens, 2004). In contrast, the marine realm is considered to have fewer obvious physical barriers, which reduces the opportunities for strict allopatry to take place (Bowen et al., 2013; Brandley et al., 2010). Most marine invertebrates reproduce via a planktonic larval stage that is followed by either a benthic or a pelagic adult phase (Thorson, 1950). This life history trait confers them with the potential to facilitate extensive gene flow through dispersal across different geographic regions (Armonies, 2001; Cowen & Sponaugle, 2009; Simkanin et al., 2019). Yet, it has been estimated (Mora et al., 2011) that 2.2 million eukaryotic species exist in the oceans, and currently described marine species represent a large fraction of the global biodiversity (Costello et al., 2017; Foggo et al., 2003). The high estimated diversity suggests dynamic diversification processes beyond the apparent scarcity of geographic barriers in the ocean.

A number of mechanisms have been proposed to account for this astonishing biodiversity and current distributions of many groups of marine organisms. Habitat choice by settling larvae has been shown to be responsible for maintaining a mosaic genetic structure in two mussel species in a small geographic scale (Barrett, 2017; Bierne et al., 2003; Frolova & Miglietta, 2020); adaptation to alternative water temperatures maintains reproductive isolation between sister species of *Halichoeres* fishes (Rocha et al., 2005); reduced gene flow between continental and oceanic islands coasts has been shown to drive divergence in shallow-water reef-associated species (Hachich et al., 2015); sister lineages of Antarctic krill species have been shown to have diverged due to the formation of circum-Antarctic water circulation and the Antarctic Polar Frontal Zone (Patarnello et al., 1996); major dispersal barriers such as the Amazonian Orinoco Plume and the Mid-Atlantic Barrier, the Isthmus of Panama, among others, are also known to promote divergence as the result of vicariance or long distance dispersal across them, followed by local adaptation (Cowman & Bellwood, 2013; Luiz et al., 2013; Varona et al., 2019). Regardless of the spatial configuration in which the divergence takes place (*i.e.* allopatric, parapatric, peripatric or sympatric), the geographic and ecological heterogeneity plays a major role in all of these mechanisms.

Neocoleoid cephalopods are marine invertebrates that comprise octopuses, cuttlefishes and squids. Many studies have dealt with systematics and macroevolution in lineages of these organisms in the last decade (Kolis & Lieberman, 2019; Lindgren et al., 2012; Strugnell et al., 2006; Strugnell & Nishiguchi, 2007; Tanner, 2018; Wani, 2011). However, very few studies have addressed diversification dynamics from a geographic perspective, covering temporal and phylogenetic hypotheses (but see (Amor et al., 2014; Brakoniecki, 1987; J. B. de L. Sales et al., 2017; Ulloa et al., 2017). Therefore, the geographic drivers of diversification in cephalopods remain poorly understood.

The increase of available molecular data for the species in the squid family Loliginidae, Lesueur, 1821, has resulted in systematic revisions with more phylogenetic accuracy. Evidence of possible cryptic species has been shown for many genera of loliginid squids(Anderson, 2000; J. B. de L. Sales et al., 2017), including three cases of potential cryptic speciation between lineages in the North and South Atlantic of the species Doryteuthis pealeii, D. pleii and Lolliguncula brevis (de Luna Sales et al., 2013; J. B. L. Sales et al., 2014). Also, a couple of studies have been published with divergence time estimates for major cephalopod lineages(Anderson, 2000; J. B. de L. Sales et al., 2017; Ulloa et al., 2017). Most species within this family are of commercial interest and their reproductive biology and ecology have been widely studied (Jereb & Roper, 2010; Martins et al., 2006; Okutani, 1995; Reid & Carstens, 2012; Vecchione, 1991). Due to their world-wide distribution (Fig. 1), and the availability of molecular and fossil data (M. Clarke & Maddock, 1988), loliginid squids represent an ideal case to study the mechanisms causing diversification in cephalopods.

**Figure 1.**
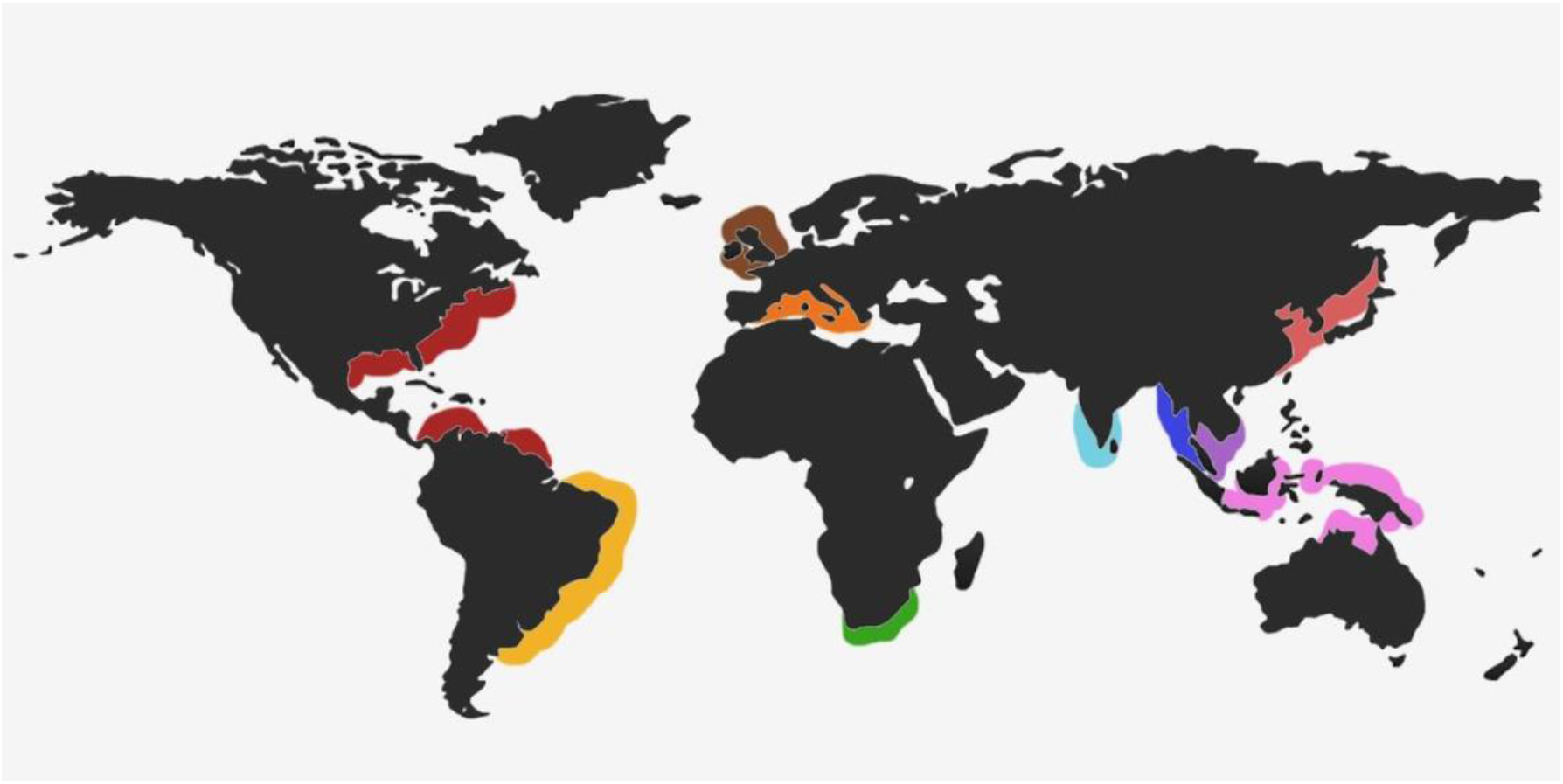
Location of the worldwide distribution of loliginid squids with assigned distributions areas to each species according to previously defined marine biogeographic regions by Spalding et al. (2007).

We aim to use a data-mining approach to estimate a dated most-inclusive phylogeny of the family Loliginidae. We coupled this with ancestral area reconstruction and ecological niche modelling to explore patterns of diversification in a spatiotemporal context. This allowed us to assess the occurrence of potential diversification mechanisms driven by geographic factors. This includes (I) isolation as a result of long-distance dispersal; (II) vicariance due to the presence of hard physical dispersal barriers; (III) divergence due to the presence of soft physical dispersal barriers (e.g. oceanic currents, river mouths); (IV) adaptation to different niches along ecological gradients.

## MATERIALS AND METHODS

### Taxon sampling and data acquisition

We used a data-mining approach to build our datasets using published sequences for 185 Operational Taxonomic Units (OTU’s) comprising 26 loliginid species. We obtained 429 sequences for three markers (167 *Cytochrome Oxidase* I, 88 *Rhodopsin* and 126 *16S*) from the Genbank database (see Table 1 for references). Sequences from *Stenoteuthis oualensis* y *Omnastrephes bartramii* were used as outgroup in all analyses.

### Phylogenetic analysis and species delimitation

Multiple sequence alignments were performed using MAFFT 7(Katoh & Standley, 2013) with the Q-INS-i strategy as implemented in the web server (http://mafft.cbrc.jp/alignment/server/) and manually edited on MESQUITE 3.04 (Maddison & Maddison, 2007). PARTITIONFINDER 1.1.1 (Lanfear et al., 2012) was used to find the best partition scheme and model of nucleotide substitution for the single locus datasets using the rcluster search algorithm and the corrected Akaike Information Criterion (AICc). The searches were performed with the following starting subsets: *COI* 1^st^, 2^nd^, 3^rd^ codon positions; *RHO* 1^st^, 2^nd^, 3^rd^ codon positions and *16S*. To construct single locus trees, Maximum Likelihood analysis were done using RAXML 8 (Stamatakis, 2014) as implemented in the CIPRES online platform (Miller et al., 2010) using 1000 bootstrap and the GTR+CAT substitution model. Moreover, we generated single locus chronograms using an uncorrelated lognormal relaxed clock in BEAST v 1.8.3(Drummond et al., 2012; Drummond & Rambaut, 2007) with default priors. We ran two independent 10,000,000 generations analyses with four chains each sampling every 1000th generation for a total sample size of 10,000 trees. Effective sample size for the estimated parameters and convergence was assessed in TRACER 1.6 (Andrew Rambaut et al., 2018). For each locus, a set of 400 trees from the estimated posterior distribution were subsampled using LOGCOMBINER 1.8.2 (A Rambaut & Drummond, 2015) after discarding 10% of the samples as burn-in. These sets of trees were finally used to infer single locus-based species delimitations using a bayesian implementation of the Generalized Mixed Yule Coalescent model (bGMYC) (Pons et al., 2006; Reid & Carstens, 2012).

### Species tree and divergence time estimation

Conflicting inter-species relationships were found with strong support (>70% bootstrap support) among phylogenies inferred from the different single-locus datasets. For this reason we decided to perform a dated species tree estimation in *BEAST (Heled & Drummond, 2010) as an alternative to concatenation. We assigned individuals to species according to the results of the previous phylogenetic analyses and used a yule tree prior for the inference. We used statoliths from the genus *Loligo* (M. R. Clarke & Fitch, 1979) to model a distribution of the node corresponding to the ancestor of the genus to calibrate the tree. For the calibration, we used a lognormal distribution with the following parameters: log(mean)= 1.64; log(standard deviation)= 0.54; and Offset= 40. Ucld.mean parameter (Uncorrelated Lognormal relaxed clock mean) was set to a uniform prior distribution with an initial value of 0.002, a maximum value of 0.003 and a lower value of 0.001. Effective sample sizes for the estimated parameters and convergence were checked in TRACER 1.6. Consensus tree was generated on TREEANNOTATOR 1.8 (A Rambaut & Drummond, 2007) using a 50% majority rule and discarding 10% as burn-in.

### Ancestral areas reconstruction

We used the Bayesian Binary MCMC and the Dispersal Extinction Cladogenesis (DEC) model as implemented in RASP (Ree & Smith, 2008; Yu et al., 2015) to obtain distribution ranges at each node. The inferred species tree was used as input for these analyses. We defined distributions areas (Fig. 1) according to previously defined marine biogeographic regions (Spalding et al., 2007) and assigned them to each species (Table S1).

### Climatic niche model approaches

The Amazonian Orinoco Plume, in the southwest Atlantic Ocean is considered as a dispersal barrier for marine organisms (Luiz et al., 2013), since the decreases in the seawater salinity in this region makes it physiologically unsuitable for most non-estuarine marine organisms. Therefore, we tested whether the (a) Amazonian Orinoco Plume is functioning as a dispersal barrier for the putative species inside *D. pleii* and *L. brevis*, and/or (b) they are currently occupying different climatic spaces thus promoting divergence. No analysis was performed for *D. pealeii* because of the lack of geographic occurrences available at databases and/or papers. For this, we used ecological niche modelling and climatic space overlapping. In both cases, we obtained georeferenced occurrence data from GBIF database in June 2017 (Global Biodiversity Information Facility, www.gbif.org) and survey records in published literature (Herke & Foltz, 2002) for the species mentioned above. The data was handled at the population level (northern and southern populations), using only the environmental information available for its region of origin (i.e. North or South Atlantic). To estimate the climatic space and the potential geographic distribution for each, we used oceanic environmental layers of Salinity, pH, Sea Surface Temperature range, Chlorophyll-A range, Dissolved Oxygen, Diffuse Attenuation mean, and Photosynthetically Available Radiation from Bio-ORACLE (Tyberghein et al., 2012). Previous eco-physiological studies have shown that these variables affect the distribution of the modeled species (Laughlin & Livingston, 1982; Martins et al., 2006; Sobrino et al., 2002; Zielinski et al., 2000). We retrieved data from the climatic layers exclusively from the West Atlantic Ocean along America. We also restricted the layers to areas with up to 1000 m (Seibel, 2007) of depth using the bathymetrical information presented by Amante and Eakins (2009).

We generated separate climatic niche models for North and South populations using data from their region of origin, and then projected them from one region to the other. The ecological niche models were built using MAXENT 3.3.3 (S. Phillips et al., 2010; S. J. Phillips & Dudík, 2008), applying the default parameters and using cross-validation to evaluate the models with 20% data. We obtained logistic models to 1 km x 1 km, with values ranging from 0 (when the area is unsuitable) to 1 (when is totally suitable) according to the climatic conditions. Finally, we used a multivariate environmental similarity surface (MESS) analysis (Elith et al., 2010) to evaluate the similarity between the North and South environments and therefore, the reliability of the projection.

We estimated the climatic space overlapping between northern and southern populations, following Broennimann and others (2012) using the *ecospat* package (Broennimann et al., 2015) as implemented in R Software (RStudio Team, 2020). The model performs a main components analysis using the environmental conditions of 10000 random locations throughout the entire study area (including both North and South regions). The climatic space estimation per population uses only the first two principal components (PC1 and PC2) and assumes a kernel distribution for the density of records to smoothen possible gaps. These gaps could be the result of sampling bias instead of conditions not being occupied by a population. The climatic space overlap is measured with the D metric (Schoener, 1970; Warren et al., 2017), ranging from 0 (when no overlap is found) to 1 (when the overlapping is complete). After that, a randomization process is used to evaluate if the estimated D value is significant according to the available conditions (Broennimann et al., 2012). A significantly low overlap between the climatic space of North and South populations could indicate climatic divergence between populations.

## RESULTS

### Phylogeny and species delimitation

The *COI* tree (Fig. 2) showed the highest resolution among the three genes, recovering the monophyly of the twenty-six loliginid species with a bootstrap >70% (except for *D. pleii*). Phylogenetic analyses based on the two other loci did not show strong support for all species, but they mostly recovered the same monophyletic groups or showed no significant conflict with the *COI* tree (Fig. S1, S2, S3). The phylogenetic analyses confirmed the previously reported geographic structure of *D. pleii, D. pealeii* and *L. brevis* (Anderson, 2000), with divergent populations in the North and South West Atlantic. This result was supported by the *COI* species delimitation analysis, in which the isolated populations resulted as distinct species (Fig. 2). Putative cryptic species or divergent lineages with geographic structure were detected inside three other clades (*Sepioteuthis lessoniana, Sepioteuthis sepioidea and Uroteuthis duvauceli*) with the Maximum Likelihood trees and the species delimitation analysis performed with *COI.*

**Figure 2.**
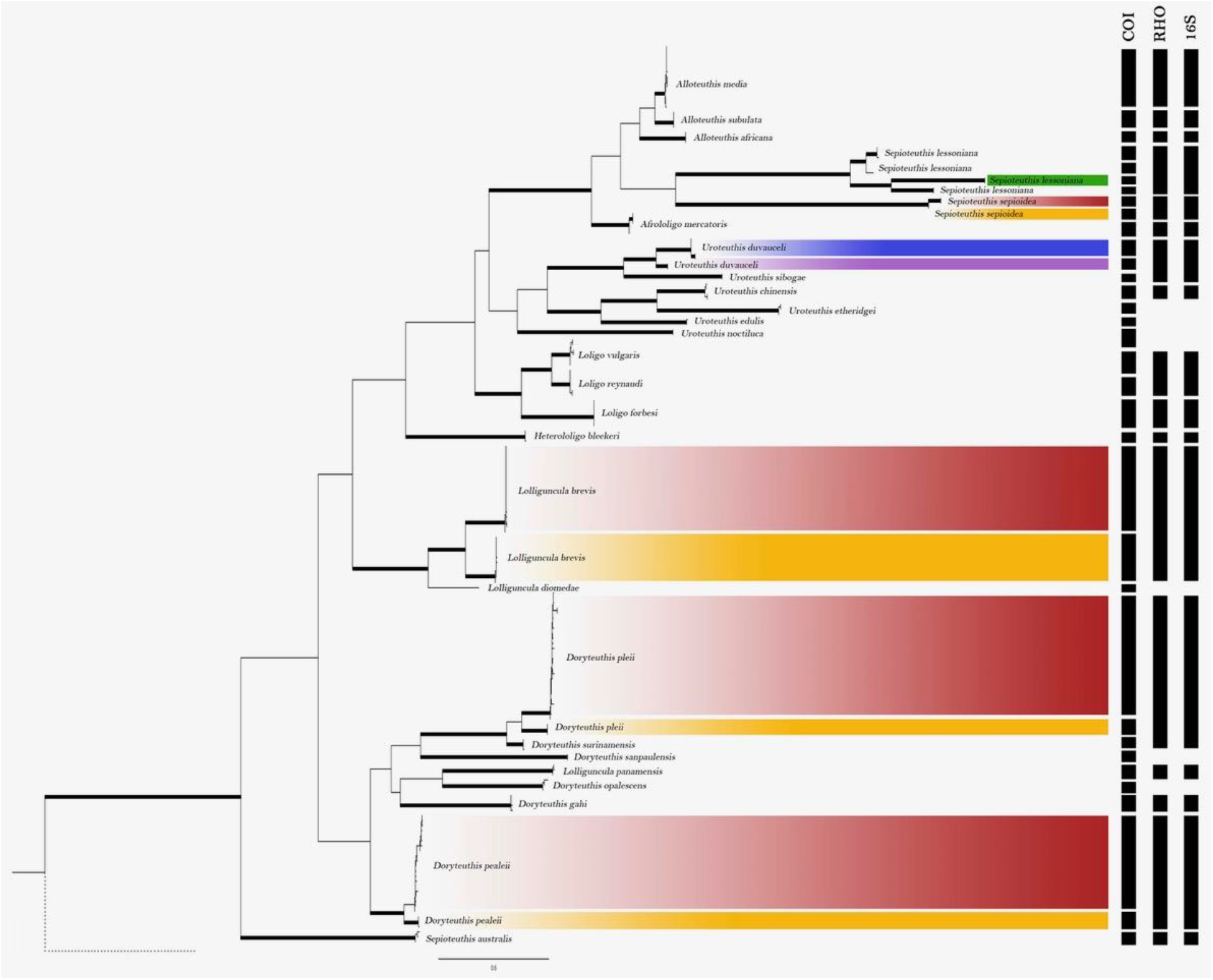
Maximum likelihood tree inferred from COI sequences. Thick branches represent bipartition with >70% bootstrap support. Bars to the right correspond to species delimitation inferred from bGMYC for each of the three loci used. Highlighted clades represent groups where cryptic speciation might have occurred and their current distribution as depicted in Figure 1.

We found incongruences regarding the relationship between species and monophyly of some genera. In order to solve the relationships between genera in the Loliginidae family, a phylogenetic analysis including all species with available sequences for the three genes was generated with a Bayesian coalescent approach (Fig. 3). This analysis clarified the evolutionary relationships between species and confirmed the monophyly from seven of the eight genera included in the study. The genus *Lolliguncula* is shown to be polyphyletic (Figure 2 and 3). This is supported by the species tree and the single locus phylogenies that show *L. panamensis* nested within the genus *Doryteuthis*.

**Figure 3.**
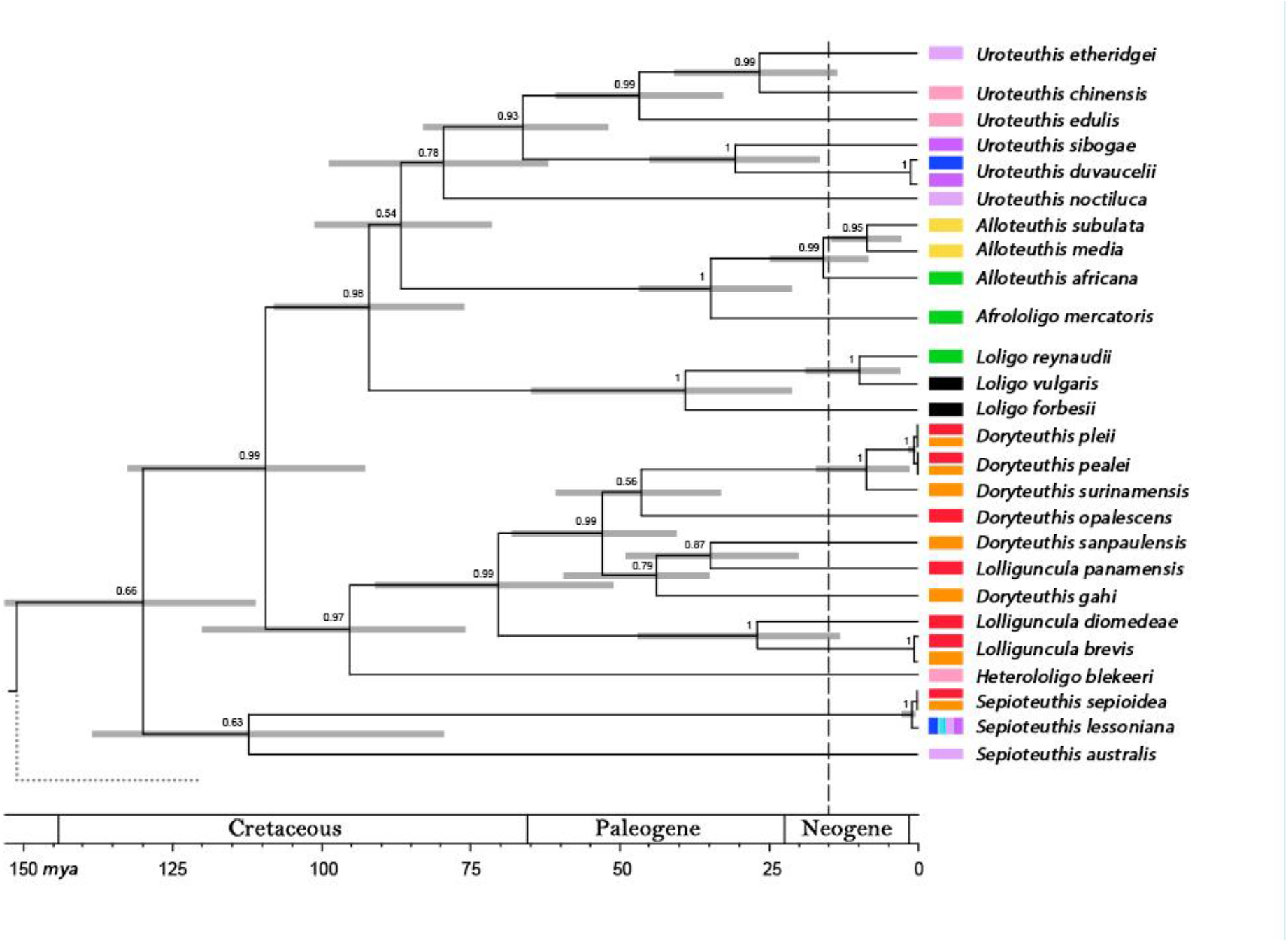
Dated species tree using information from three loci. Node bars represent 95% HPD of the node ages. Vertical discontinuous line with grey area indicated the estimated date of the closure of the Panamá isthmus. Color rectangles on the right show the current geographic distribution of each species as shown in Figure 1. Bottom scale shows geological time ages in mya.

### Ancestral area reconstruction and divergence estimation

The different models used to infer ancestral distribution ranges for nodes consistently estimated the origin and geographical dispersal of loliginid lineages (as shown in Fig. 3). A summary of the values estimated for dispersal, extinction, cladogenesis (in the DEC model), and dispersal and vicariance is presented in the Dryad Digital Repository and a description of the events is presented in the discussion.

### Climatic niche model approaches

The ecological niche models (ENM) showed low probability of suitability in the immediately adjacent area of Amazonian Orinoco Plume for both species and increased as the areas are farther from the Plume (Fig. 4). According to the projections, the South region has suitable areas for North populations from both species (Fig. 4). In contrast, the North region has suitable areas only for the southern *L. brevis* (Fig. 4d). According to the MESS analyses the environmental conditions between the origin and projections sites are highly similar except in the South to North projection, where the northernmost region of the Atlantic present high uncertainty associated (Fig. S5).

**Figure 4.**
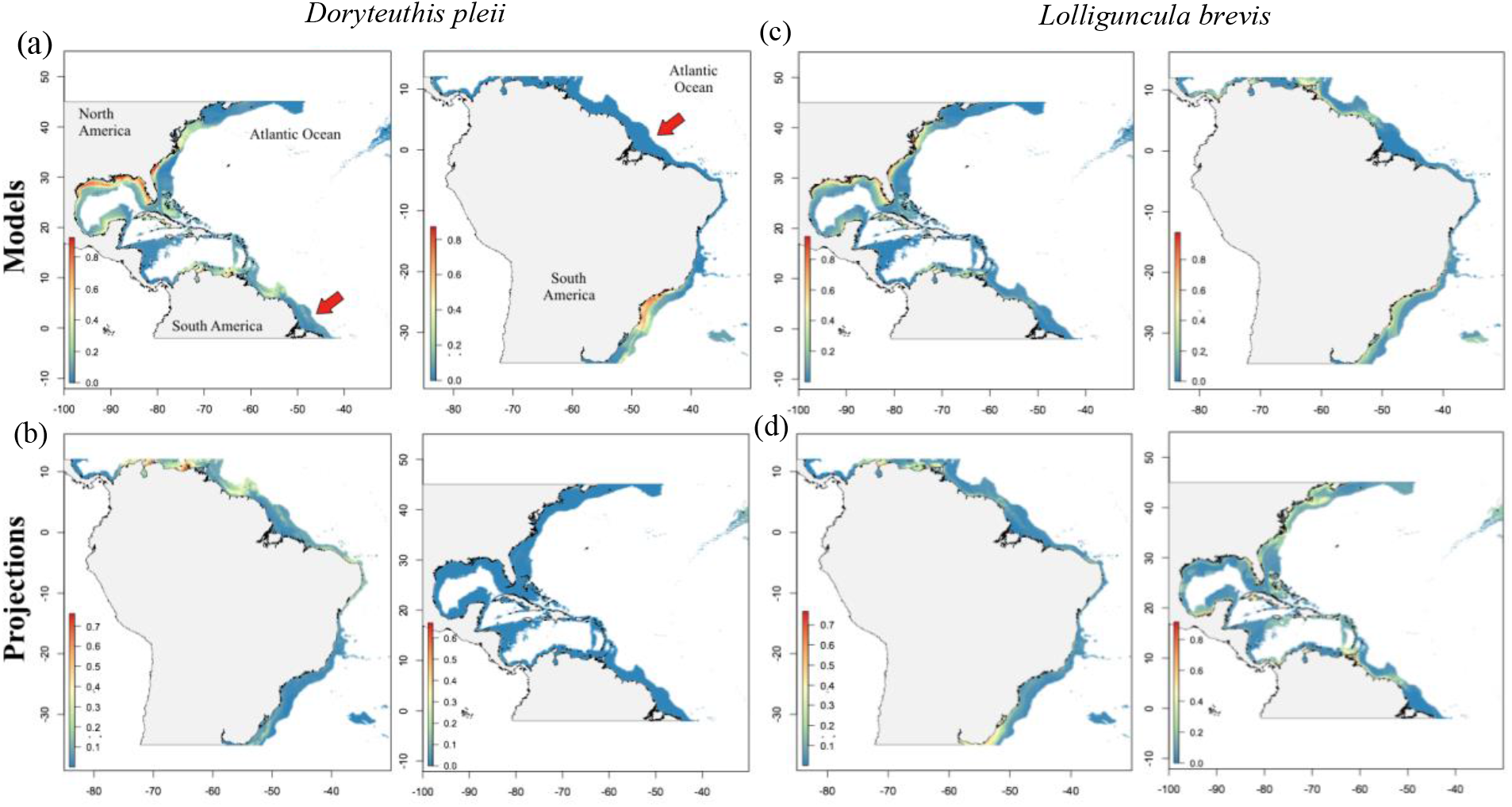
Ecological niche models for *D. pleii* and *L. brevis* and its projections. (a) and (c) show the potential geographic distribution for North and South populations of both species made by modelling the ecological niche. (b) and (d) show the projections of each model in the non-origin regions. Colors in the maps indicate the probability of suitability, with the highest probabilities in warm colors. The Amazonian Orinoco Plume region, marked with the red arrow, showed a low suitability for both species. According to the models, all populations have suitable areas in the non-origin regions, except for the South population of *D. pleii* in the North region as shown in (b).

In the climatic space overlap, we do not found evidence on climatic niche divergence between North and South populations for both species, since we found no significant overlap in the environmental space currently occupied between them (*L. brevis*: overlap=0.112, p-value=0.495; *D. pleii*: overlap=0.513, p-value=0.099).

## DISCUSSION

Having robust phylogenetic hypothesis is key to understand processes of diversification (Losos, 1996). Here we constructed a phylogeny of the family Loliginidae in order to unveil geographic drivers of diversification in these organisms. Together with ancestral areas reconstruction (AAR) and divergence time estimates among major clades, we aid the inference of the biogeographical history of the family in its early evolution. Conversely, species delimitation coupled with ecological niche model approaches revealed putative cryptic lineages that might represent recent and incipient diversification processes. The entire approach allowed us to correlate geological events and geographic heterogeneity with ancestral speciation and current distributions of extant lineages.

### New insights into phylogenetic relationships within Loliginidae

We confirmed the monophyly of all recognized species included here and clarified several between-species relationships within the family (Fig. 2 and 3). All phylogenetic analyses consistently showed the polyphyly of the genus *Lolliguncula,* as the species *L. panamensis* was nested within the *Doryteuthis* clade (Fig. 2, 3 S1, S2). The remaining two *Lolliguncula* species were monophyletic and appeared as the sister clade of the genus *Doryteuthis* (Fig. 2). The latest phylogenetic revision of *Lolliguncula* showed *L. panamensis* as the earliest divergent lineage of the genus when using *Loligo* species as outgroup (Fig. 1 and 2 in (J. B. L. Sales et al., 2014), possibly resulted from a taxon sampling artifact as no sequences of others were included in the analyses. Our results suggest a review of *L. panamensis* and plausible re-circumscription for the species to *Doryteuthis panamensis*. Moreover, previous phylogenetic analyses using both mitochondrial and nuclear markers resulted in the polyphyly of the genus *Sepioteuthis* (de Luna Sales et al., 2013). We obtained the same topology using the *COI* dataset (Fig. 2), but the species tree recovered the three species from *Sepioteuthis* as monophyletic (Fig. 2). The inferred divergence time between *S. lessoniana* and *S. sepioidea* is very recent (Fig. 2), thus, incomplete lineage sorting due to deep coalescence might explain the conflicting results found between species and gene trees. Lastly, the position of *Heterololigo bleekeri* (Ulloa et al., 2017) is confirmed with our analyses, placing this species as a sister group to the *Doryteuthis/Lolliguncula* clade with strong support (Fig. 2).

### Migration events followed by speciation characterized the early biogeographical history of the family

The results of the AAR analyses and the divergence time estimation showed that the most likely place of origin of the family is the Indo-Pacific region around 112-152 mya in the Cretaceous (Fig. 3). Migration then occurred from there in two directions: North-east and west (Supplemental Data 1). One species of the earliest divergent genus, namely *S. sepioidea,* is nowadays present in Caribbean waters. However, no strong statistical support was obtained for younger nodes that could inform the possible colonization pathway of this species from the Indo-Pacific to the Caribbean region (Fig. 3). The *Doryteuthis/Lolliguncula* clade also comprises several species currently present in the new world coasts. Inferred age and ancestral distribution of this node indicates that this ancestor could arrive to the Atlantic coast via eastward pacific migration from the Indo-Pacific and through the Atlantic long before the Panamá Isthmus closure (Fig. 3).

### Species delimitation of recent divergent populations within cryptic lineages

We found six species that comprise putative cryptic lineages (Fig. 2). Five of them had already been shown to present population differentiation in other studies (i.e., *D. pleii, D. pealei* (Herke & Foltz, 2002), *L. brevis* (de Luna Sales et al., 2013), *S. lessoniana* and *U. duvauceli* (Bergman, 2013)) while one, *Sepioteuthis sepioidea,* was detected for the first time with our phylogenetic analyses (Fig. 1). However, the bGMYC analysis supported the divergence of phylogeographic groups within these six species only on the *Cytochrome oxidase* I locus. The difference between the *COI* and *Rho* analyses could be explained by the different effective sample sizes (ESS) of mitochondrial versus nuclear genes. ESS of mitochondrial genes are smaller due to the mitochondria maternal inheritance and its haploidy, allowing loci to reach reciprocal monophyly faster (Funk & Omland, 2003; Moore, 1995). Hence, *COI* usually retrieves trees with higher resolution for recently diverged taxa. Although *16S* is also a mitochondrial gene, it has been shown that this gene is less efficient and has lower levels of taxonomic resolution (Funk & Omland, 2003), thus why the best resolution was retrieved with *COI.* However, it has been showed (J. B. de L. Sales et al., 2017) that the divergence time of the two clades within *D. pleii,* one comprising northwestern and central Caribbean Atlantic and the other southwestern Atlantic specimens, was not recent and instead about 16 million years ago. These results contradict our divergence time estimation, the latter suggesting a split between *D. plei* and *D. pealeii* no older than 5 mya. Thus, the explanation of a recent divergence between phylogeographic groups and inconsistency between molecular markers wouldn’t be supported by previous studies but could agree with our divergence time results.

### Divergence time estimates and ENMs support the Amazon-Orinoco Plume as a driver of diversification in *D. pleii* and *L. brevis*

The species *D. pleii*, *D. pealeii* and *L. brevis*. have two genetically isolated populations, each inhabiting the north and south Atlantic coast along the American continent (de Luna Sales and others, 2013). We found that the environmental conditions for the species tested (i.e., *D. pleii* and *L.brevis)* are suitable in both regions which means that northern and southern populations are occupying the same environmental space (except for the southern population of *D. pleii* in the northern Atlantic, Figure 4). However, our niche modelling projections for both species, show this area as environmentally different and unsuitable, mainly in the mouth of the Amazon, which would support the Plume as a barrier. The sea surface temperatures are strongly influenced by rainfall changes in the Amazon River basin, impacting the river discharge and consequently the sea surface salinity in the Amazon plume (Hu et al., 2004; Vizy & Cook, 2010). Low-salinity surface waters are found as far as 2000 km at an average depth of 20-30m from the mouth (Gouveia et al., 2019; Tyaquiçã et al., 2017; Varona et al., 2019). It is one of the greatest discharges of fresh water and suspended sediments in the world, previously presented as a barrier for the dispersal of other organisms such as bacterioplankton, coral and lionfish (de Souza et al., 2017; Hewson et al., 2006; Luiz et al., 2013).

Previous works on biogeography of loliginid squids (Bartol et al., 2002; de Araujo & Gasalla, 2018; Şen, 2005; Zaragoza et al., 2015) have also hypothesized that physicochemical changes in sea water triggered by events such as river drainage may serve as barriers to dispersal for adults, paralarvae and eggs in these animals. Moreover, salinity has been suggested to influence the distribution of squids, due to the narrow salinity ranges that early developmental stages can tolerate (Cinti et al., 2004).

The establishment of the Amazon River system with its current eastwards drainage to the Atlantic Ocean has been dated to occur in the mid-late Miocene, around 10.6 to 9.7 mya (Figueiredo et al., 2010). However, it was not only until the late Pliocene, around 2.5 mya, when it reached the present size and shape (Hoorn et al., 1995). Similar timeline followed the origin and establishment of the Orinoco River during the late Miocene (Cinti et al., 2004; Hoorn et al., 1995). The geological history of the rivers and their Atlantic drainage precedes and approximates our estimated divergence times of populations of all four cryptic lineages along the western Atlantic coast (i.e., *D. pleii, D. pealeii, L. brevis* and *S. sepioidea*), which resulted in splits younger than 5 mya (Figure 3). Coupled with the ecological niche modeling for *D. pleii* and *L. brevis*, these results support the hypothesis of the Amazonian Orinoco Plume as a barrier which isolated the species in two populations and prevented posterior gene flow, resulting in their consequent divergence.

### Concluding remarks

Our study provides a robust insight into the historical biogeography of the Loliginidae family. Our phylogenetic analysis coupled with the ancestral area reconstruction confirmed that the origin of this family highly likely occurred in the Indo-Pacific region around 112-152 mya, in the Cretaceous. Additionally, we were able to confirm the monophyly of all currently recognized species, to reveal the placement of Lolliguncula panamensis within the Doryteuthis clade and to find what seems to be a possible new cryptic lineage (i.e. *Sepioteuthis sepioidea)*. Further on, our divergence time estimates and ecological niche analyses uncovered the mechanisms and evolutionary history behind the genetic isolation of *D. pleii* and *L. brevi*s populations, with the Amazon-Orinoco Plume playing a very relevant role in the diversification of these species and possibly the other ones distributed along the western Atlantic coast. Our results encourage further studies of the impact that the increase of such barrier may have not only on squids but also in other species.

## DATA AVAILABILITY STATEMENT

All data used in this study was downloaded from open sources as listed in the Methods section and the Supplementary Table 1. The supporting data generated here (i.e. alignments, raw phylogenetic trees, coordinates, layers, occurrences, R script) are available from the Dryad Digital Repository

## BIOSKETCH

Gabrielle Genty has a masters in science from the University of St. Andrews, Scotland, and is currently working as a research field assistant for Flinders University in Australia. Her main interests are in historical biogeography and evolutionary genetics of marine organisms.

## Author contribution

C.J.P.H., E.A.R. and G.G. conceived the project; and all authors analyzed the data, worked on the figures and wrote the manuscript.

## Acknowledgements

We would like to thank Daniel Cadena and Andrew J. Crawford for their suggestions and guidance during the early stages of this investigation.

