## Supplementary tables and figures for "Geographic drivers of diversification in loliginid squids with an emphasis on the western Atlantic species"

**Table S1.** GenBank accession numbers, sampling location (*i.e.* locality) and geographic region/s (as defined for the Ancestral Areas Reconstruction analysis) of the sequences used in the phylogenetic analyses.

| Genus | Species | Locality | Geographic Region/AAR | References | *COI* | *16S* | *Rhodopsin* |
|---|---|---|---|---|---|---|---|
| Ommastrephes | Ommastrephes bartramii | Unknown | Unknown/ ABCDEFG | Strugnell et al., 2005*; Carlini and Graves, 1999**; Takumiya et al.*** | AF000057** | AB191132*** | AY616915* |
| Sthenoteuthis | Sthenoteuthis oualaniensis | Unknown | Unknown/ HGF | Bonnaud et al., 1994*; Carlini and Graves, 1999**; Strugnell et al., 2004*** | AF000069** | X79582* | AY545185*** |
| Afrololigo | Afrololigo mercatoris | South Africa | Benguela Current/C | Anderson, 2000 | AF075390 | AF110085 | |
| Afrololigo | Afrololigo mercatoris | South Africa | Benguela Current/C | Anderson, 2000a | AF075390 | AF110085 | |
| Afrololigo | Afrololigo mercatoris | South Africa | Benguela Current/C | Anderson et al., 2008 | EU668101 | | EU668060 |
| Alloteuthis | Afrololigo mercatoris | South Africa | Benguela Current/C | Anderson, 2000*; Anderson et al., 2008** | AF075390* | AF110085* | EU668060** |
| Alloteuthis | Alloteuthis africana | Angola | Benguela Current/C | Anderson et al., 2008 | EU668061 | EU668102 | EU668026 |
| Alloteuthis | Alloteuthis africana | Angola | Benguela Current/C | Anderson et al., 2008 | EU668070 | EU668111 | EU668032 |
| Alloteuthis | Alloteuthis africana | Angola | Benguela Current/C | Anderson et al., 2008 | EU668069 | EU668110 | EU668033 |
| Alloteuthis | Alloteuthis media | Mola di Bali, Italy | Mediterranean Sea/A | Anderson et al., 2008 | EU668088 | EU668129 | EU668048 |
| Alloteuthis | Alloteuthis media | Galicia, España | Iberian Coastal/A | Anderson et al., 2008 | EU668083 | EU668124 | EU668043 |

| | | | | | | | |
|---|---|---|---|---|---|---|---|
| Alloteuthis | Alloteuthis media | France: Bay of Seine (English Channel, eastern Atlantic Ocean) | Celtic Biscay Shelf/A | Anderson et al., 2008 | EU668097 | EU668138 | EU668056 |
| Alloteuthis | Alloteuthis media | France: Banyuls-sur-Mer (western Mediterranean) | Mediterranean Sea/A | Anderson et al., 2008 | EU668096 | EU668137 | EU668055 |
| Alloteuthis | Alloteuthis media | France: Banyuls-sur-Mer (western Mediterranean) | Mediterranean Sea/A | Anderson et al., 2008 | EU668095 | EU668136 | EU668054 |
| Alloteuthis | Alloteuthis media | Italy: Mola di Bari (Adriatic Sea) | Mediterranean Sea/A | Anderson et al., 2008 | EU668094 | EU668135 | EU668053 |
| Alloteuthis | Alloteuthis media | Italy: Mola di Bari (Adriatic Sea) | Mediterranean Sea/A | Anderson et al., 2008 | EU668093 | EU668134 | EU668052 |
| Alloteuthis | Alloteuthis media | Italy: Mola di Bari (Adriatic Sea) | Mediterranean Sea/A | Anderson et al., 2008 | EU668092 | EU668133 | EU668051 |
| Alloteuthis | Alloteuthis media | Italy: Mola di Bari (Adriatic Sea) | Mediterranean Sea/A | Anderson et al., 2008 | EU668091 | EU668132 | |
| Alloteuthis | Alloteuthis media | Italy: Mola di Bari (Adriatic Sea) | Mediterranean Sea/A | Anderson et al., 2008 | EU668090 | EU668131 | EU668050 |
| Alloteuthis | Alloteuthis media | Italy: Mola di Bari (Adriatic Sea) | Mediterranean Sea/A | Anderson et al., 2008 | EU668089 | EU668130 | EU668049 |
| Alloteuthis | Alloteuthis media | Spain: Galicia | Iberian Coastal/A | Anderson et al., 2008 | EU668088 | EU668129 | EU668048 |
| Alloteuthis | Alloteuthis media | Spain: Galicia | Iberian Coastal/A | Anderson et al., 2008 | EU668087 | EU668128 | EU668047 |

| | | | | | | | |
|---|---|---|---|---|---|---|---|
| Alloteuthis | Alloteuthis media | Spain: Galicia | Iberian Coastal/A | Anderson et al., 2008 | EU668086 | EU668127 | EU668046 |
| Alloteuthis | Alloteuthis media | Spain: Galicia | Iberian Coastal/A | Anderson et al., 2008 | EU668085 | EU668126 | EU668045 |
| Alloteuthis | Alloteuthis subulata | Mola di Bali, Italy | Mediterranean Sea/A | Anderson et al., 2008 | EU668098 | EU668139 | EU668057 |
| Alloteuthis | Alloteuthis subulata | Italy: Mola di Bari (Adriatic Sea) | Mediterranean Sea/A | Anderson et al., 2008 | EU668100 | EU668141 | EU668059 |
| Alloteuthis | Alloteuthis subulata | Italy: Mola di Bari (Adriatic Sea) | Mediterranean Sea/A | Anderson et al., 2008 | EU668099 | EU668140 | EU668058 |
| Alloteuthis | Alloteuthis subulata | Italy: Mola di Bari (Adriatic Sea) | Mediterranean Sea/A | Anderson et al., 2008 | EU668098 | EU668139 | EU668057 |
| Doryteuthis | Doryteuthis gahi | Falkland Islands | Patagonian Shelf/D | Anderson, 2000 | AF075399 | AF110076 | |
| Doryteuthis | Doryteuthis gahi | Falkland Islands | Patagonian Shelf/D | Sales et al., 2013 | KF854068 | KF854030 | KF854106 |
| Doryteuthis | Doryteuthis gahi | Falkland Islands | Patagonian Shelf/D | Sales et al., 2013 | KF854067 | KF854029 | KF854105 |
| Doryteuthis | Doryteuthis gahi | Falkland Islands | Patagonian Shelf/D | Sales et al., 2013 | KF854066 | KF854028 | KF854104 |
| Doryteuthis | Doryteuthis opalescens | Monterey, CA, USA | California Current/F | Anderson, 2000 | AF075395 | AF110077 | |
| Doryteuthis | Doryteuthis opalescens | California, USA | California Current/F | Sales et al., 2013 | KF854070 | KF854032 | KF854108 |
| Doryteuthis | Doryteuthis opalescens | California, USA | California Current/F | Sales et al., 2013 | KF854069 | KF854031 | KF854107 |
| Doryteuthis | Doryteuthis pealei | Maine | Northeast US Continental Shelf/B | Anderson, 2000 | AF075408 | | |

| | | | | | | |
|---|---|---|---|---|---|---|
| Doryteuthis | Doryteuthis pealei | Gulf de Mexico | Gulf de Mexico/E | Herke and Foltz, 2002 | AF207916 | |
| Doryteuthis | Doryteuthis pealei | North Western Atlantic, USA | North America/B | Herke and Foltz, 2002 | AF207924 | |
| Doryteuthis | Doryteuthis pealei | Rhode Island, USA | Northeast US Continental Shelf/B | Anderson, 2000 | AF075391 | AF110079 |
| Doryteuthis | Doryteuthis pealei | Gulf de Mexico | Gulf de Mexico/E | Herke and Foltz, 2002 | AF207926 | |
| Doryteuthis | Doryteuthis pealei | North Western Atlantic, USA | North America/B | Herke and Foltz, 2002 | AF207925 | |
| Doryteuthis | Doryteuthis pealei | Gulf de Mexico | Gulf de Mexico/E | Herke and Foltz, 2002 | AF207923 | |
| Doryteuthis | Doryteuthis pealei | Gulf de Mexico | Gulf de Mexico/E | Herke and Foltz, 2002 | AF207922 | |
| Doryteuthis | Doryteuthis pealei | Gulf de Mexico | Gulf de Mexico/E | Herke and Foltz, 2002 | AF207921 | |
| Doryteuthis | Doryteuthis pealei | North Western Atlantic, USA | North America/B | Herke and Foltz, 2002 | AF207920 | |
| Doryteuthis | Doryteuthis pealei | North Western Atlantic, USA | North America/B | Herke and Foltz, 2002 | AF207919 | |
| Doryteuthis | Doryteuthis pealei | Gulf de Mexico | Gulf de Mexico/E | Herke and Foltz, 2002 | AF207918 | |
| Doryteuthis | Doryteuthis pealei | North Western Atlantic, USA | North America/B | Herke and Foltz, 2002 | AF207917 | |
| Doryteuthis | Doryteuthis pealei | North Western Atlantic, USA | North America/B | Herke and Foltz, 2002 | AF207915 | |
| Doryteuthis | Doryteuthis pealei | Gulf de Mexico | Gulf de Mexico/E | Herke and Foltz, 2002 | AF207914 | |
| Doryteuthis | Doryteuthis pealei | North Western Atlantic, USA | North America/B | Herke and Foltz, 2002 | AF207913 | |
| Doryteuthis | Doryteuthis pealei | Gulf de Mexico | Gulf de Mexico/E | Herke and Foltz, 2002 | AF207912 | |

| | | | | | | | |
|---|---|---|---|---|---|---|---|
| Doryteuthis | Doryteuthis pealei | North Western Atlantic, USA | North America/B | Herke and Foltz, 2002 | AF207911 | | |
| Doryteuthis | Doryteuthis pealei | Gulf de Mexico | Gulf de Mexico/E | Herke and Foltz, 2002 | AF207910 | | |
| Doryteuthis | Doryteuthis pealei | North Western Atlantic, USA | North America/B | Herke and Foltz, 2002 | AF207909 | | |
| Doryteuthis | Doryteuthis pealei | Rhode Island, USA | Northeast US Continental Shelf/B | Sales et al., 2013 | KF854064 | KF854027 | KF854103 |
| Doryteuthis | Doryteuthis pealei | Rhode Island, USA | Northeast US Continental Shelf/B | Sales et al., 2013 | KF854065 | KF854026 | KF854102 |
| Doryteuthis | Doryteuthis pealei | Bragança, Pará State | Brazil/D | Sales et al., 2013 | KF854055 | KF854017 | KF854093 |
| Doryteuthis | Doryteuthis pealei | Bragança, Pará State | Brazil/D | Sales et al., 2013 | KF854054 | KF854016 | KF854092 |
| Doryteuthis | Doryteuthis pealei | Bragança, Pará State | Brazil/D | Sales et al., 2013 | KF854053 | KF854015 | KF854091 |
| Doryteuthis | Doryteuthis pleii | Unknown | Unknown | Anderson, 2000 | AF075404 | AF110080 | |
| Doryteuthis | Doryteuthis pleii | Gulf de Mexico | Gulf de Mexico/E | Herke and Foltz, 2002 | AF207947 | | |
| Doryteuthis | Doryteuthis pleii | North Western Atlantic, USA | North America/B | Herke and Foltz, 2002 | AF207946 | | |
| Doryteuthis | Doryteuthis pleii | Gulf de Mexico | Gulf de Mexico/E | Herke and Foltz, 2002 | AF207945 | | |
| Doryteuthis | Doryteuthis pleii | North Western Atlantic, USA | North America/B | Herke and Foltz, 2002 | AF207944 | | |
| Doryteuthis | Doryteuthis pleii | Gulf de Mexico | Gulf de Mexico/E | Herke and Foltz, 2002 | AF207943 | | |
| Doryteuthis | Doryteuthis pleii | North Western Atlantic, USA | North America/B | Herke and Foltz, 2002 | AF207942 | | |
| Doryteuthis | Doryteuthis pleii | Gulf de Mexico | Gulf de Mexico/E | Herke and Foltz, 2002 | AF207941 | | |

| | | | | | | | |
|---|---|---|---|---|---|---|---|
| Doryteuthis | Doryteuthis pleii | Gulf de Mexico | Gulf de Mexico/E | Herke and Foltz, 2002 | AF207940 | | |
| Doryteuthis | Doryteuthis pleii | North Western Atlantic, USA | North America/B | Herke and Foltz, 2002 | AF207939 | | |
| Doryteuthis | Doryteuthis pleii | Gulf de Mexico | Gulf de Mexico/E | Herke and Foltz, 2002 | AF207938 | | |
| Doryteuthis | Doryteuthis pleii | Gulf de Mexico | Gulf de Mexico/E | Herke and Foltz, 2002 | AF207937 | | |
| Doryteuthis | Doryteuthis pleii | North Western Atlantic, USA | North America/B | Herke and Foltz, 2002 | AF207936 | | |
| Doryteuthis | Doryteuthis pleii | Gulf de Mexico | Gulf de Mexico/E | Herke and Foltz, 2002 | AF207935 | | |
| Doryteuthis | Doryteuthis pleii | North Western Atlantic, USA | North America/B | Herke and Foltz, 2002 | AF207934 | | |
| Doryteuthis | Doryteuthis pleii | North Western Atlantic, USA | North America/B | Herke and Foltz, 2002 | AF207933 | | |
| Doryteuthis | Doryteuthis pleii | Gulf de Mexico | Gulf de Mexico/E | Herke and Foltz, 2002 | AF207932 | | |
| Doryteuthis | Doryteuthis pleii | North Western Atlantic, USA | North America/B | Herke and Foltz, 2002 | AF207931 | | |
| Doryteuthis | Doryteuthis pleii | North Western Atlantic, USA | North America/B | Herke and Foltz, 2002 | AF207930 | | |
| Doryteuthis | Doryteuthis pleii | North Western Atlantic, USA | North America/B | Herke and Foltz, 2002 | AF207929 | | |
| Doryteuthis | Doryteuthis pleii | Gulf de Mexico | Gulf de Mexico/E | Herke and Foltz, 2002 | AF207928 | | |
| Doryteuthis | Doryteuthis pleii | North Western Atlantic, USA | North America/B | Herke and Foltz, 2002 | AF207927 | | |
| Doryteuthis | Doryteuthis pleii | Salinas, Para State | Brazil/D | Sales et al., 2013 | KF854050 | KF854012 | KF854088 |
| Doryteuthis | Doryteuthis pleii | Baia Da Traição, Paraiba State | Brazil/D | Sales et al., 2013 | KF854051 | KF854013 | KF854089 |

| | | | | | | | |
|---|---|---|---|---|---|---|---|
| Doryteuthis | Doryteuthis pleii | Iracema, Santa Catarina State | Brazil/D | Sales et al., 2013 | KF854052 | KF854014 | KF854090 |
| Doryteuthis | Doryteuthis pleii | Gulf de Mexico | Gulf de Mexico/E | Sales et al., 2013 | KF854063 | KF854025 | KF8540101 |
| Doryteuthis | Doryteuthis pleii | Gulf de Mexico | Gulf de Mexico/E | Sales et al., 2013 | KF854062 | KF854024 | KF8540100 |
| Doryteuthis | Doryteuthis pleii | Gulf de Mexico | Gulf de Mexico/E | Sales et al., 2013 | KF854061 | KF854023 | KF8540099 |
| Doryteuthis | Doryteuthis sanpaulensis | Rio Grande, Rio Grande Do Sul State | Brazil/D | Sales et al., 2013 | KF854060 | KF854022 | KF854098 |
| Doryteuthis | Doryteuthis sanpaulensis | Rio Grande, Rio Grande Do Sul State | Brazil/D | Sales et al., 2013 | KF854059 | KF854021 | KF854097 |
| Doryteuthis | Doryteuthis surinamensis | Cabo Norte, Amapá State | Brazil/D | Sales et al., 2013 | KF854058 | KF854020 | KF854096 |
| Doryteuthis | Doryteuthis surinamensis | Cabo Norte, Amapá State | Brazil/D | Sales et al., 2013 | KF854057 | KF854019 | KF854095 |
| Doryteuthis | Doryteuthis surinamensis | Cabo Norte, Amapá State | Brazil/D | Sales et al., 2013 | KF854056 | KF854018 | KF854094 |
| Heterololigo | Heterololigo bleekeri | Aomori, Honshu, Japan | Oyashio Current/G | Anderson, 2000a | AF075388 | AF110074 | |
| Heterololigo | Heterololigo bleekeri | Kingo Ito, Japan | Japan/G | Sales et al., 2013 | KF854072 | KF854034 | KF854110 |
| Heterololigo | Heterololigo bleekeri | Kingo Ito, Japan | Japan/G | Sales et al., 2013 | KF854071 | KF854033 | KF854109 |
| Loligo | Loligo forbesii | Plymouth, UK; * unknown | Celtic Biscay Shelf/A | Anderson, 2000a*; Strugnell et al. 2004** | AF075402 | AF110075 | AY545184 |
| Loligo | Loligo forbesii | West Coast of Scotland | Celtic Biscay Shelf/A | Sales et al., 2013 | KF854077 | KF854039 | KF854115 |

| | | | | | | | |
|---|---|---|---|---|---|---|---|
| Loligo | Loligo forbesii | West Coast of Scotland | Celtic Biscay Shelf/A | Sales et al., 2013 | KF854078 | KF854040 | KF854116 |
| Loligo | Loligo reynaudii | South Africa | Benguela Current/C | Anderson, 2000 | AF075406 | AF110081 | |
| Loligo | Loligo reynaudii | Tsirsirkana, South Africa | South Africa/C | Sales et al., 2013 | KF854073 | KF854035 | KF854111 |
| Loligo | Loligo reynaudii | Tsirsirkana, South Africa | South Africa/C | Sales et al., 2013 | KF854074 | KF854036 | KF854112 |
| Loligo | Loligo vulgaris | Plymouth, U.K. | Celtic Biscay Shelf/A | Anderson, 2000 | AF075397 | AF110082 | |
| Loligo | Loligo vulgaris | Plymouth, U.K. | Celtic Biscay Shelf/A | Anderson, 2000a | AF075397 | AF110082 | |
| Loligo | Loligo vulgaris | Lisbon, Portugal | Iberian Coastal/A | Sales et al., 2013 | KF854075 | KF854037 | KF854113 |
| Loligo | Loligo vulgaris | Lisbon, Portugal | Iberian Coastal/A | Sales et al., 2013 | KF854076 | KF854038 | KF854114 |
| Lolliguncula | Lolliguncula brevis | Galveston, Texas, USA, Gulf de Mexico | Gulf de Mexico/E | Anderson, 2000a*; Strugnell et al., 2005** | AF075396 | AF110084* | AY616916** |
| Lolliguncula | Lolliguncula brevis | Pena Island near Salinas, Pará state, Brazil | North Brazil Shelf/D | Sales et al., 2014 | KF266740 | KF266728 | KF266752 |
| Lolliguncula | Lolliguncula brevis | Pena Island near Salinas, Pará state, Brazil | North Brazil Shelf/D | Sales et al., 2014 | KF266741 | KF266729 | KF266753 |
| Lolliguncula | Lolliguncula brevis | Pena Island near Salinas, Pará state, Brazil | North Brazil Shelf/D | Sales et al., 2014 | KF266742 | KF266730 | KF266754 |
| Lolliguncula | Lolliguncula brevis | Pena Island near Salinas, Pará state, Brazil | North Brazil Shelf/D | Sales et al., 2014 | KF273944 | KF273939 | KF273953 |

| | | | | | | | |
|---|---|---|---|---|---|---|---|
| Lolliguncula | Lolliguncula brevis | Pena Island near Salinas, Pará state, Brazil | North Brazil Shelf/D | Sales et al., 2014 | KF273945 | KF273938 | KF273954 |
| Lolliguncula | Lolliguncula brevis | Pena Island near Salinas, Pará state, Brazil | North Brazil Shelf/D | Sales et al., 2014 | KF273946 | KF273935 | KF273955 |
| Lolliguncula | Lolliguncula brevis | Pena Island near Salinas, Pará state, Brazil | North Brazil Shelf/D | Sales et al., 2014 | KF273947 | KF273940 | KF273956 |
| Lolliguncula | Lolliguncula brevis | Pena Island near Salinas, Pará state, Brazil | North Brazil Shelf/D | Sales et al., 2014 | KF273948 | KF273937 | KF273957 |
| Lolliguncula | Lolliguncula brevis | Pena Island near Salinas, Pará state, Brazil | North Brazil Shelf/D | Sales et al., 2014 | KF273949 | KF273936 | KF273958 |
| Lolliguncula | Lolliguncula brevis | Pena Island near Salinas, Pará state, Brazil | North Brazil Shelf/D | Sales et al., 2014 | KF273950 | KF273934 | KF273959 |
| Lolliguncula | Lolliguncula brevis | Baia da Traição, Paraíba state, Brazil | East Brazil Shelf/D | Sales et al., 2014 | KF266737 | KF266725 | KF266749 |
| Lolliguncula | Lolliguncula brevis | Baia da Traição, Paraíba state, Brazil | East Brazil Shelf/D | Sales et al., 2014 | KF266738 | KF266726 | KF266750 |
| Lolliguncula | Lolliguncula brevis | Baia da Traição, Paraíba state, Brazil | East Brazil Shelf/D | Sales et al., 2014 | KF266739 | KF266727 | KF266751 |
| Lolliguncula | Lolliguncula brevis | Jequié, Bahia state, Brazil | East Brazil Shelf/D | Sales et al., 2014 | KF266736 | KF266724 | KF266748 |
| Lolliguncula | Lolliguncula brevis | Jequié, Bahia state, Brazil | East Brazil Shelf/D | Sales et al., 2014 | KF273943 | KF273933 | KF273952 |
| Lolliguncula | Lolliguncula brevis | Guaiabim Beach near Valença, Bahia state, Brazil | East Brazil Shelf/D | Sales et al., 2014 | KF266743 | KF266731 | KF266755 |

| | | | | | | | |
|---|---|---|---|---|---|---|---|
| Lolliguncula | Lolliguncula brevis | Grauçá Beach near Caravelas, Bahia state, Brazil | East Brazil Shelf/D | Sales et al., 2014 | KF273942 | KF273941 | KF273951 |
| Lolliguncula | Lolliguncula brevis | Grauçá Beach near Caravelas, Bahia state, Brazil | East Brazil Shelf/D | Sales et al., 2014 | KF266735 | KF266723 | KF266747 |
| Lolliguncula | Lolliguncula brevis | Ciudad del Carmen,Southern Gulf de Mexico | Gulf de Mexico/E | Sales et al., 2014 | KF854136 | KF854126 | KF854145 |
| Lolliguncula | Lolliguncula brevis | Ciudad del Carmen,Southern Gulf de Mexico | Gulf de Mexico/E | Sales et al., 2014 | KF854137 | KF854127 | |
| Lolliguncula | Lolliguncula brevis | Ciudad del Carmen,Southern Gulf de Mexico | Gulf de Mexico/E | Sales et al., 2014 | | KF854128 | |
| Lolliguncula | Lolliguncula brevis | Ciudad del Carmen,Southern Gulf de Mexico | Gulf de Mexico/E | Sales et al., 2014 | KF854138 | KF854129 | KF854146 |
| Lolliguncula | Lolliguncula brevis | Ciudad del Carmen,Southern Gulf de Mexico | Gulf de Mexico/E | Sales et al., 2014 | KF854139 | KF854130 | KF854147 |
| Lolliguncula | Lolliguncula brevis | Ciudad del Carmen,Southern Gulf de Mexico | Gulf de Mexico/E | Sales et al., 2014 | KF854140 | KF854131 | KF854148 |
| Lolliguncula | Lolliguncula brevis | Ciudad del Carmen,Southern Gulf de Mexico | Gulf de Mexico/E | Sales et al., 2014 | KF854141 | KF854132 | KF854149 |

| | | | | | | | |
|---|---|---|---|---|---|---|---|
| Lolliguncula | Lolliguncula brevis | Ciudad del Carmen,Southern Gulf de Mexico | Gulf de Mexico/E | Sales et al., 2014 | KF854142 | KF854133 | |
| Lolliguncula | Lolliguncula brevis | Ciudad del Carmen,Southern Gulf de Mexico | Gulf de Mexico/E | Sales et al., 2014 | KF854143 | KF854134 | KF854150 |
| Lolliguncula | Lolliguncula brevis | Ciudad del Carmen,Southern Gulf de Mexico | Gulf de Mexico/E | Sales et al., 2014 | KF854144 | KF854135 | KF854151 |
| Lolliguncula | Lolliguncula diomedeae | East Tropical Pacific, México | Pacific Central American Coastal/F | Lindgren, 2010 | EU735357 | EU735243 | |
| Lolliguncula | Lolliguncula panamensis | Bahia Las Animas, Gulf of California, México | Gulf de Mexico/E | Sales et al., 2014 | KF266744 | KF266732 | KF266756 |
| Lolliguncula | Lolliguncula panamensis | Bahia Las Animas, Gulf of California, México | Gulf de Mexico/E | Sales et al., 2014 | KF266745 | KF266733 | KF266757 |
| Lolliguncula | Lolliguncula panamensis | Bahia Las Animas, Gulf of California, México | Gulf de Mexico/E | Sales et al., 2014 | KF266746 | KF266734 | KF266758 |
| Sepioteuthis | Sepioteuthis australis | South Australia | Australia/H | Anderson, 2000 | AF075401 | AF110087 | |
| Sepioteuthis | Sepioteuthis australis | Otago, New Zealand | New Zealand Shelf/H | Anderson, 2000 | AF075386 | AF110086 | |
| Sepioteuthis | Sepioteuthis australis | Australia | Australia/H | Carline and Graves, 1999; Strugnell et al., 2005* | AF000065 | | AY616917* |

| | | | | | | | |
|---|---|---|---|---|---|---|---|
| Sepioteuthis | Sepioteuthis lessoniana | Sulawesi, Indonesia | Indonesian Sea/H | Anderson, 2000 | AF075405 | AF110088 | |
| Sepioteuthis | Sepioteuthis lessoniana | Green Island, Moreton Bay, Australia | East Centrtal Australian Shelf/H | Anderson, 2000 | AF075393 | | |
| Sepioteuthis | Sepioteuthis lessoniana | Unknown | Unknown/HG | Shiao et al. | AY131050 | AY131020 | |
| Sepioteuthis | Sepioteuthis lessoniana | Unknown | Unknown/HG | Shiao et al. | AY131055 | AY131021 | |
| Sepioteuthis | Sepioteuthis lessoniana | Unknown | Unknown/HG | Shiao et al. | AY131056 | AY131023 | |
| Sepioteuthis | Sepioteuthis lessoniana | Unknown | Unknown/HG | Shiao et al. | AY131064 | AY131027 | |
| Sepioteuthis | Sepioteuthis lessoniana | Unknown | Unknown/HG | Shiao et al. | AY131065 | AY131028 | |
| Sepioteuthis | Sepioteuthis lessoniana | Sulawesi, Indonesia | Indonesian Sea/H | Anderson, 2000a | AF075405 | AF110089 | |
| Sepioteuthis | Sepioteuthis lessoniana | Unknown | Unknown/HG | Liu et al.*; Shiao et al.**;Strugnell et al., 2005*** | AY131036** | AJ001649* | AY616918*** |
| Sepioteuthis | Sepioteuthis lessoniana | Durban, South Africa | South Africa/H | Sales et al., 2013 | KF854086 | KF854048 | KF854124 |
| Sepioteuthis | Sepioteuthis lessoniana | Durban, South Africa | South Africa/H | Sales et al., 2013 | KF854087 | KF854049 | KF854125 |
| Sepioteuthis | Sepioteuthis sepioidea | Bahamas | Caribbean island/E | Anderson, 2000 | AF075392 | AF110090 | |
| Sepioteuthis | Sepioteuthis sepioidea | Bahamas | Caribbean island/E | Anderson, 2000a | AF075392 | AF110090 | |
| Sepioteuthis | Sepioteuthis sepioidea | Barra Grande, Bahia State | Brazil/D | Sales et al., 2013 | KF854085 | KF854047 | KF854123 |
| Uroteuthis | Uroteuthis chinensis | Gulf of Thailand | Gulf of Thailand/H | Anderson, 2000 | AF075394 | AF110091 | |
| Uroteuthis | Uroteuthis chinensis | Unknown | Unknown | Liu et al. | | AJ000105 | |

| | | | | | | | |
|---|---|---|---|---|---|---|---|
| Uroteuthis | Uroteuthis chinensis | Xiamen, China | South China Sea/G | Sin et al., 2009 | EU349439 | EU349478 | |
| Uroteuthis | Uroteuthis chinensis | Terengganu, Malaysia | Malaysia/H | Sales et al., 2013 | KF854079 | KF854041 | KF854117 |
| Uroteuthis | Uroteuthis duvauceli | Andaman Sea | Bay of Bengal/H | Anderson, 2000 | AF075400 | AF110092 | |
| Uroteuthis | Uroteuthis duvauceli | Gulf of Thailand | Gulf of Thailand/H | Anderson, 2000a | EU349465 | EU349492 | |
| Uroteuthis | Uroteuthis duvauceli | Hong Kong, China | South China Sea/G | Sin et al., 2009 | EU349463 | EU349490 | |
| Uroteuthis | Uroteuthis duvauceli | Yamaguchi, Japón | East Sea of Japan/H | Anderson, 2000a | AF075400 | AF110092 | |
| Uroteuthis | Uroteuthis duvauceli | Terengganu, Malaysia | Malaysia/H | Sales et al., 2013 | KF854084 | KF854046 | KF854122 |
| Uroteuthis | Uroteuthis duvauceli | Terengganu, Malaysia | Malaysia/H | Sales et al., 2013 | KF854083 | KF854045 | KF854121 |
| Uroteuthis | Uroteuthis duvauceli | Terengganu, Malaysia | Malaysia/H | Sales et al., 2013 | KF854082 | KF854044 | KF854120 |
| Uroteuthis | Uroteuthis edulis | Shangai, China | East China Sea/G | Sin et al., 2009 | EU349462 | EU349488 | |
| Uroteuthis | Uroteuthis edulis | Shangai, China | East China Sea/G | Sin et al., 2009 | EU349456 | EU349485 | |
| Uroteuthis | Uroteuthis etheridgei | Moreton Bay, Australia | East Central Australian Shelf/H | Anderson, 2000 | AF075389 | AF110094 | |
| Uroteuthis | Uroteuthis etheridgei | Moreton Bay, Australia | East Central Australian Shelf/H | Anderson, 2000a | AF075389 | AF110094 | |
| Uroteuthis | Uroteuthis noctiluca | Australia | Australia/H | Anderson, 2000 | AF075403 | AF110095 | |
| Uroteuthis | Uroteuthis noctiluca | Australia | Australia/H | Anderson, 2000a | AF075403 | AF110095 | |

| | | | | | | | |
|---|---|---|---|---|---|---|---|
| Uroteuthis | Uroteuthis sibogae | Terengganu, Malaysia | Malaysia/H | Sales et al., 2013 | KF854081 | KF854043 | KF854119 |
| Uroteuthis | Uroteuthis sibogae | Terengganu, Malaysia | Malaysia/H | Sales et al., 2013 | KF854080 | KF854042 | KF854118 |
| Uroteuthis | Uroteuthis sp. | Melbourne, Australia | Southeast Australian Shelf/H | Strugnell et al., 2005 | AY616889 | AY616881 | AY616919 |

**Table S2.** Overlapping of climatic niche among the north and south populations of *Lolliguncula brevis* and *Doryteuthis pleii*. As higher are the D values, higher is the overlapping. The equivalency test evaluates if the climatic space of the two populations are identical, whereas the similarity test evaluates if the climatic space of the two populations are more or less similar (according to D value) than expect by chance. Because all p-values are higher than 0.05, we did not find evidence about the climatic space occupied by the populations be more or less similar than expected only by chance.

| **Population 1** | **Population 2** | **Overlap (D)** | **Equivalency test (p-value)** | **Similarity test (p-value)** |
|---|---|---|---|---|
| *L. brevis* - North | *L. brevis* - South | 0.112012166 | 1 | 0.495049505 |
| *L. brevis* - North | *D. pleii* - North | 0.052062744 | 1 | 0.465346535 |
| *L. brevis* - South | *D. pleii* - North | 0.284161683 | 1 | 0.108910891 |
| *L. brevis* - North | *D. pleii* - South | 0.038875399 | 1 | 0.524752475 |
| *L. brevis* - South | *D. pleii* - South | 0.335345722 | 0.909090909 | 0.128712871 |
| *D. pleii* - North | *D. pleii* - South | 0.512622431 | 0.909090909 | 0.099009901 |

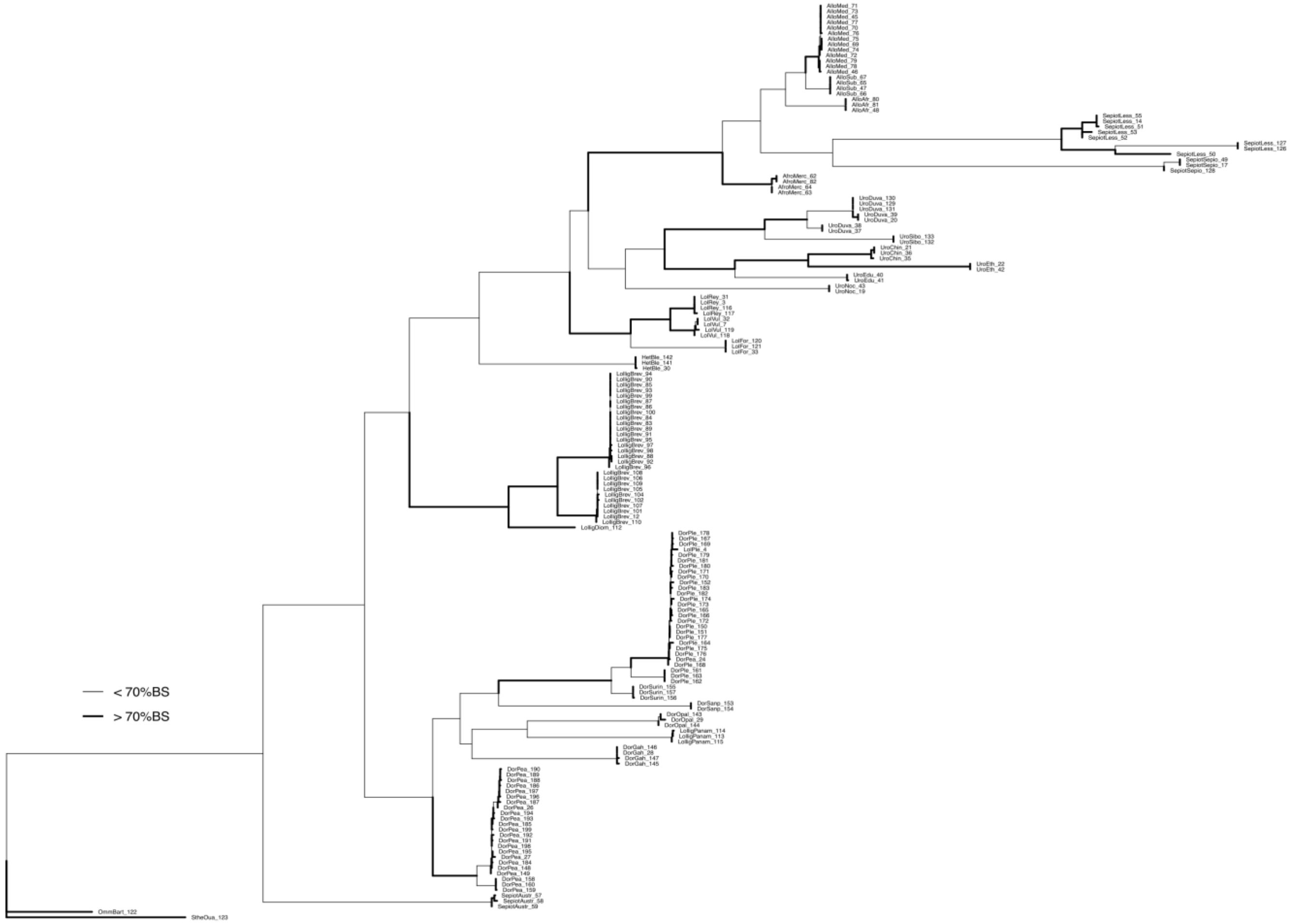

**Fig S1.** ***Cytochrome c oxidase subunit I*** **(COI) Maximum Likelihood tree.** Thicker branches represent bootstrap supports over the 70% threshold. Tree file is also available in the Dryad repository (link).

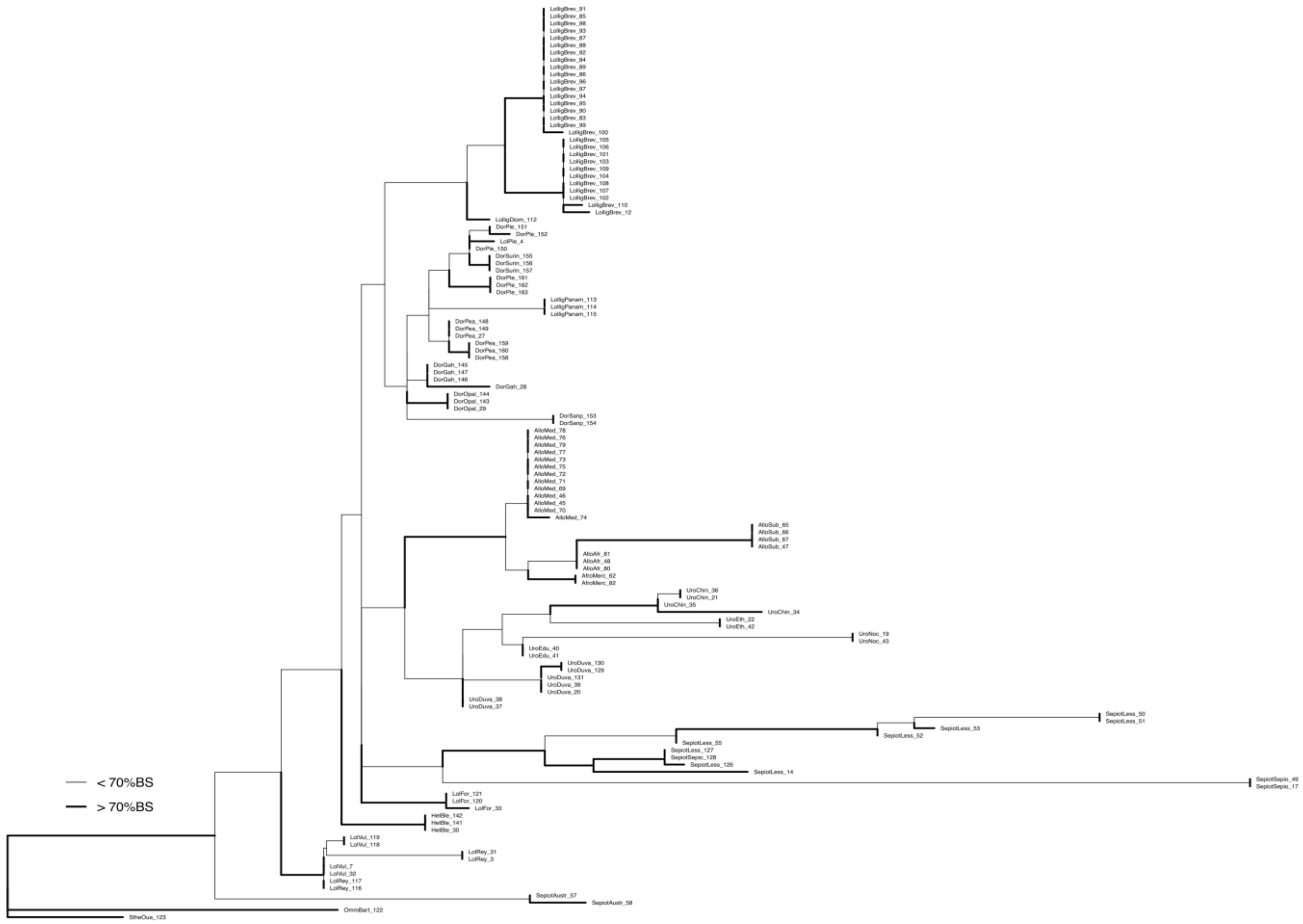

**Fig S2. 16S Maximum Likelihood tree.** Thicker branches represent bootstrap supports over the 70% threshold. Tree file is also available in the Dryad repository (link).

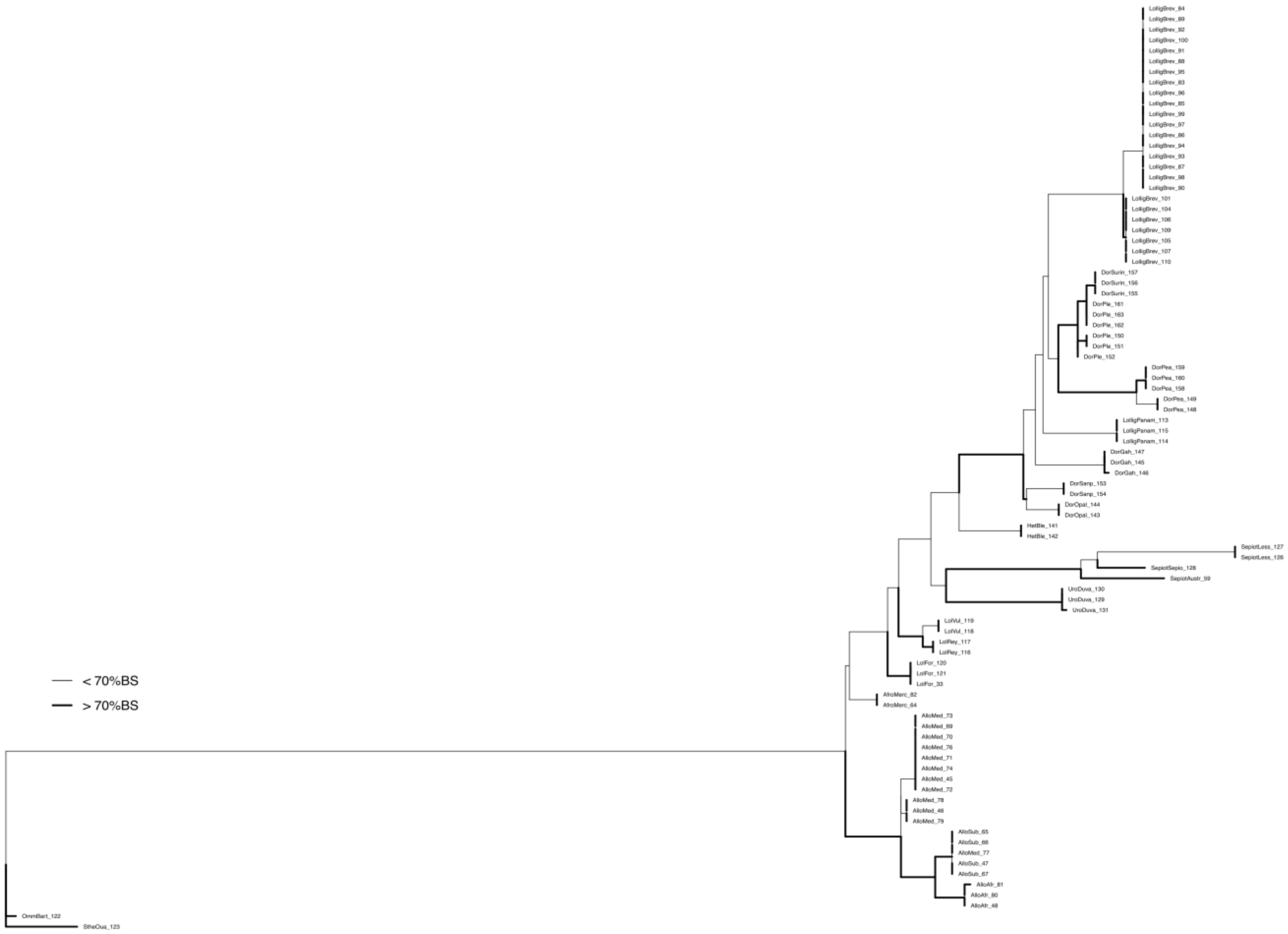

**Fig S3.** ***Rhodopsin*** **(Rho) Maximum Likelihood tree.** Thicker branches represent bootstrap supports over the 70% threshold. Tree file is also available in the Dryad repository (link).

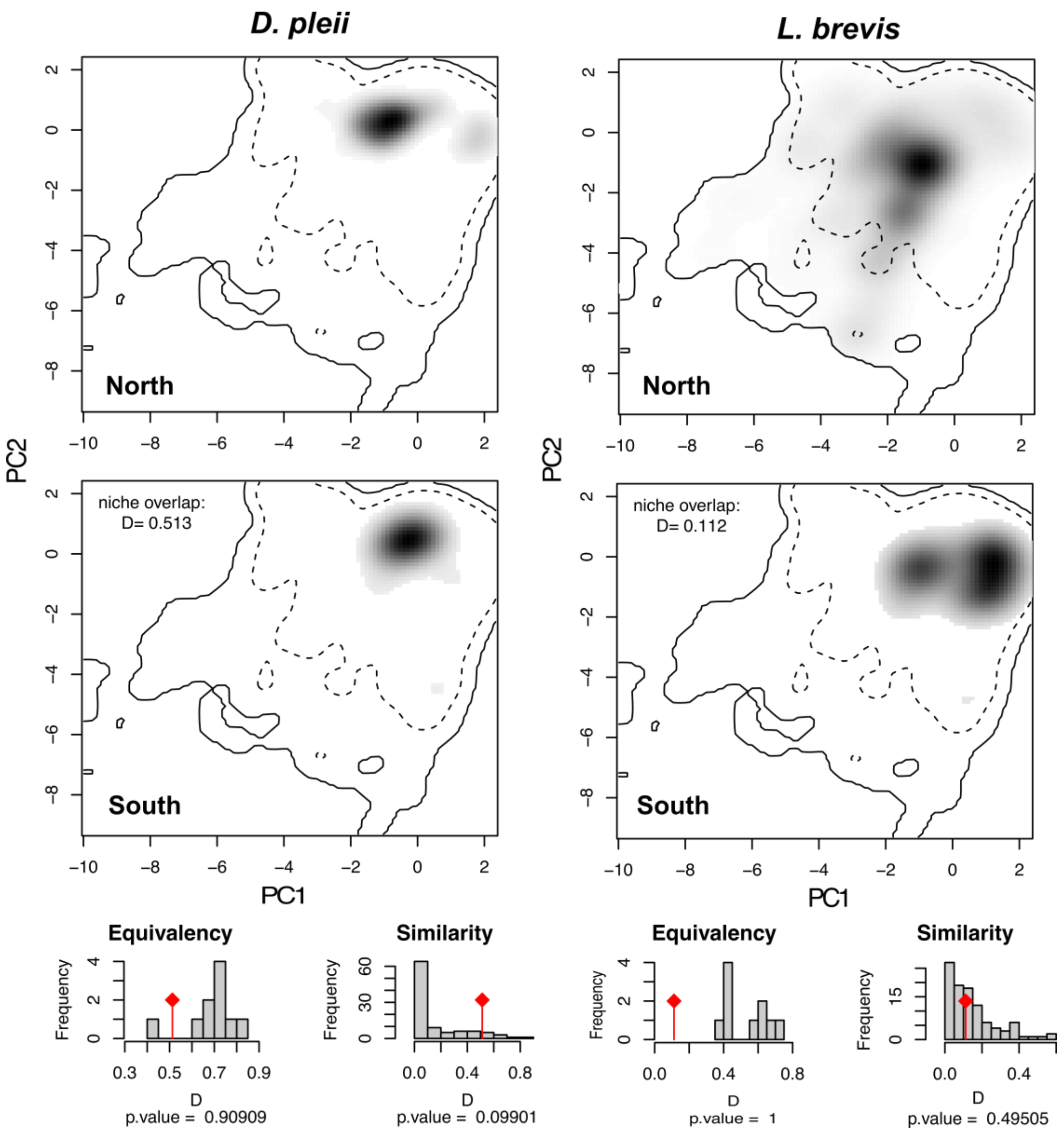

**Fig S4.** Climatic space occupied by the north and south populations from *Doryteuthis pleii* (left panel) and, north and south populations from *Lolliguncula brevis* (right panel). The solid and dashed lines indicate the 100% and 95% of the available space respectively, along the two first axes of the PCA of the environmental variables. The black pixels show the space occupied by each population, with darker pixels indicating a higher density of occurrences. In the lower panel is show the null model to estimate the significance of the equivalency and similarity tests, with the red rhombus specifying observed D values.

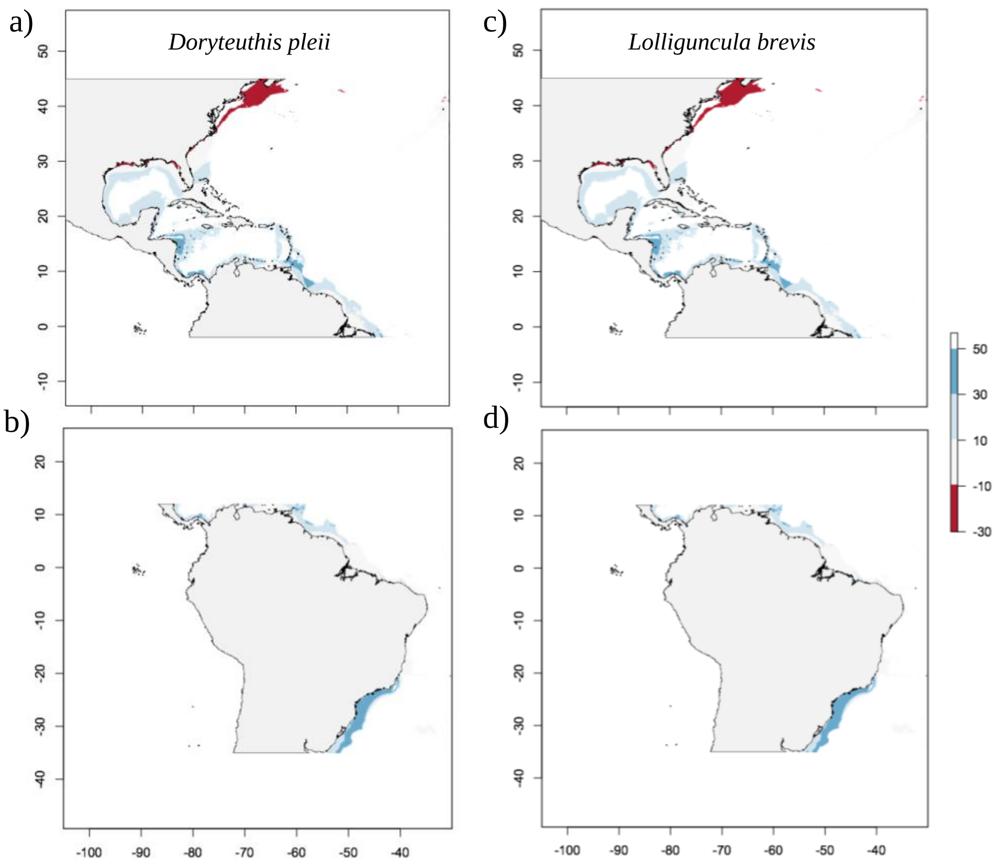

**Fig S5.** Results of MESS analyses, showing the similarity of environmental conditions between the regions where the models were generated and the region where the models were projected: (a) and (c) North America in regard to South America; (b) and (d) South America in regard to North America. The colors scale indicates the similarity, from blue (high similarity) to red (low similarity), being the latter, the areas with highest uncertainty.
